# Exploring genetic diversity in the species-rich genus *Liolaemu*s: The interplay among isolation by distance, body size, and environmental variability

**DOI:** 10.1101/2025.01.21.634185

**Authors:** M. R. Ruiz-Monachesi, F. B. Cruz, J. J. Martínez

## Abstract

Intraspecific genetic variation enhances the specieś capacity to endure diverse environments. Such variation may drive genetic divergence among populations, often manifesting as isolation by distance (IBD). Here, we investigated how ecological, environmental, life-history, morphological, and phylogenetic factors, shape IBD and genetic diversity (π) in *Liolaemus* lizards. Further we explored differences related to reproductive modes and taxonomic groups. Using data from GenBank, we examined two mitochondrial genes: Cytb (86 species) and 12S rRNA (37 species). We integrated geographic information for each species to evaluate genetic differentiation concerning spatial distribution. We calculated the IBD slope (β_IBD_), Manteĺs *r*, and π for each gene, then tested for an association between β_IBD_ and π. Phylogenetic multiple linear regression revealed that IBD, in the case of Cytb, was present in 56.16% of species and was negatively associated with snout-vent length (SVL). At the same time β_IBD-Cytb_ was positively linked to geographic range size. Additionally, π of Cytb gene declined with increasing temperature range variability and body size. In contrast, for 12S rRNA, IBD was present in 42.42% of species but showed no relation with SVL; instead, phylogenetic history played a more substantial role in determining π-12S rRNA. Remarkably, only Cytb showed a positive association between β and π. These results underscore different evolutionary patterns across the two genes and suggest that both, isolation by distance, alongside thermal heterogeneity, shape genetic diversity within particular reproductive modes and phylogenetic contexts in *Liolaemus*.

## Introduction

Genetic variation within and among populations, underpins the genetic structure of species (Frankham et al., 2002; Maya-Garcia et al. 2017). This variability is essential for persistence in heterogeneous and rapidly changing environments, enhancing the adaptive potential of populations (Frankham, 1995; Jamieson and Allendorf, 2012; Ekroth et al., 2019). Such variation is generated and maintained across various microevolutionary processes, such as random mutations, genetic drift, and natural or sexual selection (Lande, 1976; Lynch et al., 2016). Additionally, the spatial distribution of populations can influence genetic variation; when populations become geographically isolated, gene flow diminishes, leading to genetic divergence and, in some cases, isolation by distance (IBD). Genetic similarity is expected to decrease as geographic distance increases and then, separate populations may present isolation by distance (Wright, 1943; Hancock and Hendrick, 2018). Though, IBD patterns often arise in species with limited dispersal capabilities and they may be less pronounced or absent in species with high dispersal potential (Hancock and Hendrick, 2018). Various factors have been proposed as predictors of IBD patterns. These include ecological and behavioural traits (e.g., diet—herbivore vs. carnivore—, foraging habits —sit and wait vs. wide forager), life-history attributes (e.g., reproductive strategies, pace-of-life syndrome), taxonomic group, ontogenetic stage (e.g., adults vs. larvae in amphibians), morphological features (e.g., limb length). Body size, for instance, can influence dispersal ability and habitat use, where larger species often occupying broader ranges and potentially show weaker IBD patterns than smaller and more restricted species.

Understanding the determinants of intraspecific genetic diversity is equally critical for conservation and evolutionary biology. Species at greater extinction risk often exhibit reduced genetic diversity due to small, isolated populations susceptible to genetic drift and inbreeding (Lande, 1976; Hedrick and Kalinowski, 2000; Frankham, 2005; Rivers et al., 2014; O’Leary et al., 2015). Effective population size (N_e_) is a key determinant; larger populations generally maintain higher genetic diversity, buffering against genetic drift and promoting gene flow. Body size has negatively correlated with genetic diversity in various taxa, such as mammals, bees, and butterflies (Brüniche-Olsen et al., 2018; López-Uribe et al., 2019; Mackintosh et al., 2019). Moreover, life history traits can shape genetic diversity (Romiguier et al., 2014; Ellegren and Galtier 2016), with species investing heavily in parental care and longevity often exhibiting lower diversity than short-lived, high-fecundity ones (Romiguier et al., 2014). Natural history characteristics, such as antipredator strategies, can predict genetic diversity too; for example, aposematic (colourful) frogs often have high genetic diversity (Medina et al., 2024). Ecological contexts, such as habitat size or type, may also shape genetic diversity. Freshwater species typically exhibit lower genetic diversity than marine species (Ward et al., 1994), and genetic diversity in some fish increases with habitat size (Bolnick and Ballare, 2020). Extrinsic factors such as geographic area size (mainland vs. island distribution) and climatic factors such as, precipitation, air temperature and latitudinal gradients, were also linked to variation in genetic diversity (Miraldo et al., 2016; Smith et al., 2017; Brüniche-Olsen et al., 2018; Pelletier and Carstens, 2018; Amador et al., 2024; Navas Martínez et al., 2024). Finally, phylogeny and the type of genetic marker studied (coding vs. noncoding loci) can further influence genetic diversity among different organisms (Romiguier et al., 2014; Barrow et al., 2021; Segovia-Ramiréz et al., 2023). Thus, various factors interact in complex and taxon-specific ways, making it challenging to reach a broad consensus about the determinants of intraspecific genetic diversity.

Reptiles, the third most endangered tetrapod group, are increasingly at risk of extinction (Sinervo et al., 2010; Böhm et al., 2013; IUCN red list). Within reptiles, lizards often show IBD patterns (Jenkins et al., 2010); though, these patterns do not necessarily drive an increase in the speciation rates in Squamate reptiles (Singhal et al., 2022a). Most IBD patterns studies in lizards have employed biogeographical and phylogenetic approaches (e.g., Howes et al., 2006; Hurston et al., 2009; Díaz-Cardenas et al., 2017; Reilly et al., 2024) or have focused on niche-related issues (Fenker et al., 2024), behavioural variation (Keren-Rotem et al., 2024) or landscape genetics (Fonseca et al., 2024; Vasconcelos et al., 2024). Regarding the study of determinants of genetic diversity in reptiles, different factors were proposed as explainers of this diversity as: the number of museum occurrence records (a proxy for sampling effort), geographic position within a species’ range and climatic/environmental factors (Singhal et al., 2017, 2022 b; Grundler et al., 2019; Rivera et al., 2020) the geographic range size (Lau et al., 2024) or the size of settlements (Wall et al., 2024). Also, environmental heterogeneity, climatic and ecological environmental factors can promote genetic diversity in reptiles (Rivera et al., 2020; Vasconcelos et al., 2024). Moreover, the relative importance of range size, reproductive mode, and phylogeny, may vary depending on whether mitochondrial or nuclear genes are examined (Larkin et al., 2023).

The diverse and extensively studied genus *Liolaemus* (Abdala et al., 2021) contitutes an ideal model to further explore these relationships. With 290 species (Uetz et al., 2024), distributed from central Peru to Tierra del Fuego (Argentina and Chile) and elevation ranging from sea level to 5400 m (Cerdeña et al., 2021), these lizards occupy a wide variety of environment such as forest, halophyte, rocky, sandy, and terrestrial environments (Abdala et al., 2021). Also, present variable use of habitat (arboreal specialists, sand-dweller, saxicolous, and terrestrial generalists; Tulli et al, 2011, 2012; Abdala et al., 2021) and diet type (herbivorous, insectivorous and omnivorous; Espinoza et al., 2004; Ocampo et al., 2024). They are divided into two major phylogenetic clades: the Argentinean group (*Eulaemus*) and the Chilean group (*Liolaemus sensu stricto*; Laurent, 1983) and include species with oviparous and viviparous reproductive modes (Schulte et al., 2000; Esquerre et al., 2019). Evidence for IBD patterns in *Liolaemus* present opposite trends, some species, like *Liolaemus tenuis* and *L. lemniscatus*, exhibit clear IBD patterns (Vera-Escalona, 2012; Cab-Sulub and Álvarez-Castañeda, 2022), while others, such as *L. wiegmanni*, does not (Villamil et al., 2017). Mixed results have also been reported in species such as *L. bibronii* and *L. gracilis* (Morando et al., 2007). Genetic diversity studies in *Liolaemus* have largely focused on taxonomic, phylogenetic, and phylogeographic questions (Morando et al., 2007; Olave et al., 2011; Camargo et al., 2012; Grummer et al., 2018), with fewer studies emphasizing evolutionary and population genetics perspectives (e.g., Muñoz-Mendoza et al., 2017). Notably, no research has yet examined, the association between IBD and body size or in broad-scale determinants of genetic diversity in an evolutionary context for this genus.

In this study, we address three main objectives: (1) Estimate the slope (β_IBD_) of genetic differentiation vs. geographic distance for each species to assess potential determinants of the magnitude of IBD. We also test for body size correlation with the presence of IBD, hypothesizing that larger species may exhibit weaker IBD patterns (Brüniche-Olsen et al., 2018). (2) Estimate species-specific nucleotide diversity (π) and identify potential drivers, including morphological, life-history, and climatic factors. We also consider the role of environmental heterogeneity contrasting species with different reproductive modes (oviparous vs. viviparous) and phylogenetic groupings (*Eulaemus* vs. *Liolaemus sensu stricto*). (3) Investigate whether there is a positive association between β_IBD_ and π-values. We expect that species showing stronger IBD patterns might also present higher nucleotide diversity (Vasconcelos et al., 2024). By integrating these approaches, we aim to elucidate if ecological, morphological, life-history, phylogenetic, and environmental factors shape genetic diversity and spatial genetic structure in this species-rich genus.

## Material and Methods

### DNA sequence acquisition

We performed a comprehensive search of the GenBank database (National Center for Biotechnology Information) up to October 24, 2023, to gather all available DNA sequences for *Liolaemus* lizards. For each sequence, we recorded the species name (or a tentative taxonomic assignment whether the identification was uncertain), GenBank accession and GI numbers, specimen identifiers, sequence length (bp), DNA molecule type, and associated publication details (authors, study title, and journal) when were available (details provided in Supplementary S1). Each referenced study was reviewed for data accuracy and reliability.

### Data filtration and preparation

Initial filtering excluded records that mentioned *Liolaemus* but not provide relevant genetic sequences. We focused on the most studied markers, including only species with at least five different sequences to ensure an adequate representation of genetic variation within each species (Barrow et al., 2021). To avoid pseudo replication, we ensured to select sequences from distinct vouchers specimens. Taxonomic identities were verified using multiple systematic references (Supplementary S2). This process addressed ongoing taxonomic revision, ensured accurate species assignments and accounted for species complexes that had been split or renamed (e.g., *Liolaemus bibronii*, *L. elongatus*, *L. kingii*, *L. nigroviridis*, *L. wiegmanni*; see Supplementary S2). Only sequences linked to known geographic coordinates were retained for further subsequent analyses. After evaluating various genetic markers, we focused on Cytb and 12S rRNA, as they were the most widely represented in the dataset (Fig. 1).

**Fig.1:**
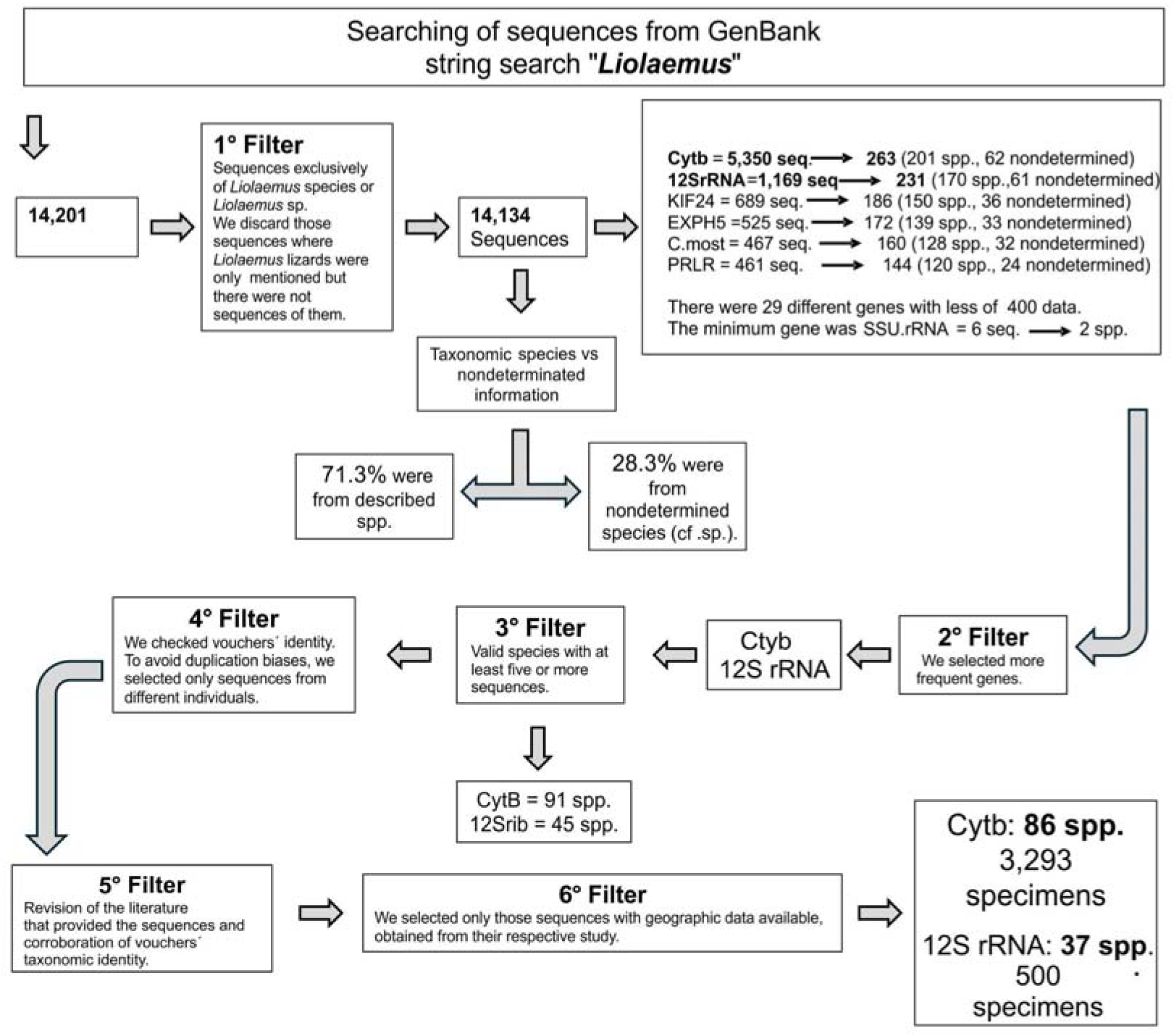
Flow diagram that illustrates the main steps used to assemble the final datasets of *Liolaemus* lizard sequences from the GenBank. Six successive filters were applied to the initial search results to ensure data accuracy and completeness, resulting in the final datasets for Cytb and 12S rRNA analyses.

### Metrics of genetic variation

All analyses were performed in R version 4.3.3 (R Developed Core Team 2024). We downloaded sequences using the “read.GenBank”, saving them as FASTA files, with the “write.dna” (both functions from *ape* package; Paradis and Schliep, 2019). Alignments were conducted for each species separately using MUSCLE v.3.8.31 (Edgar, 2004) within SeaView5 v.5.0.5 (Supplementary S3) and subsequently edited in BioEdit sequences editor v.7.1 (Hall, 1999).

To calculate nucleotide diversity (π) per species, aligned-edited FASTA files were converted into DNA sequences format using the “readAAStringSet” (*msa* package; Bodenhofer et al., 2015) and “as.DNAbin” (*ape* package; Paradis and Schliep 2019) functions. We then calculated π using the “nuc.div” from the *pegas* package (Paradis, 2010). This function sums the number of differences between pairs of sequences divided by the number of comparisons.

### Phylogenetic tree construction

Recognizing that species share a common phylogenetic history and should not be treated as independent statistical units (Felsenstein, 1985), we performed phylogenetic corrections in our analyses. We constructed two meta-trees: one for species with Cytb data and one for those with 12S rRNA, using Mesquite v2.74 (Maddison and Maddison, 2017). These trees integrated systematic information from multiple studies (Supplementary S2). Branch lengths were standardised to one, and subsequently scaled to be ultrametric using “compute.brlen” (*ape* package; Paradis and Schliep 2019). Trees visualization was done with “plotTree” (*phytools* package; Revell, 2024), resulting in the two phylogenetic frameworks used here (Fig.2).

**Fig. 2:**
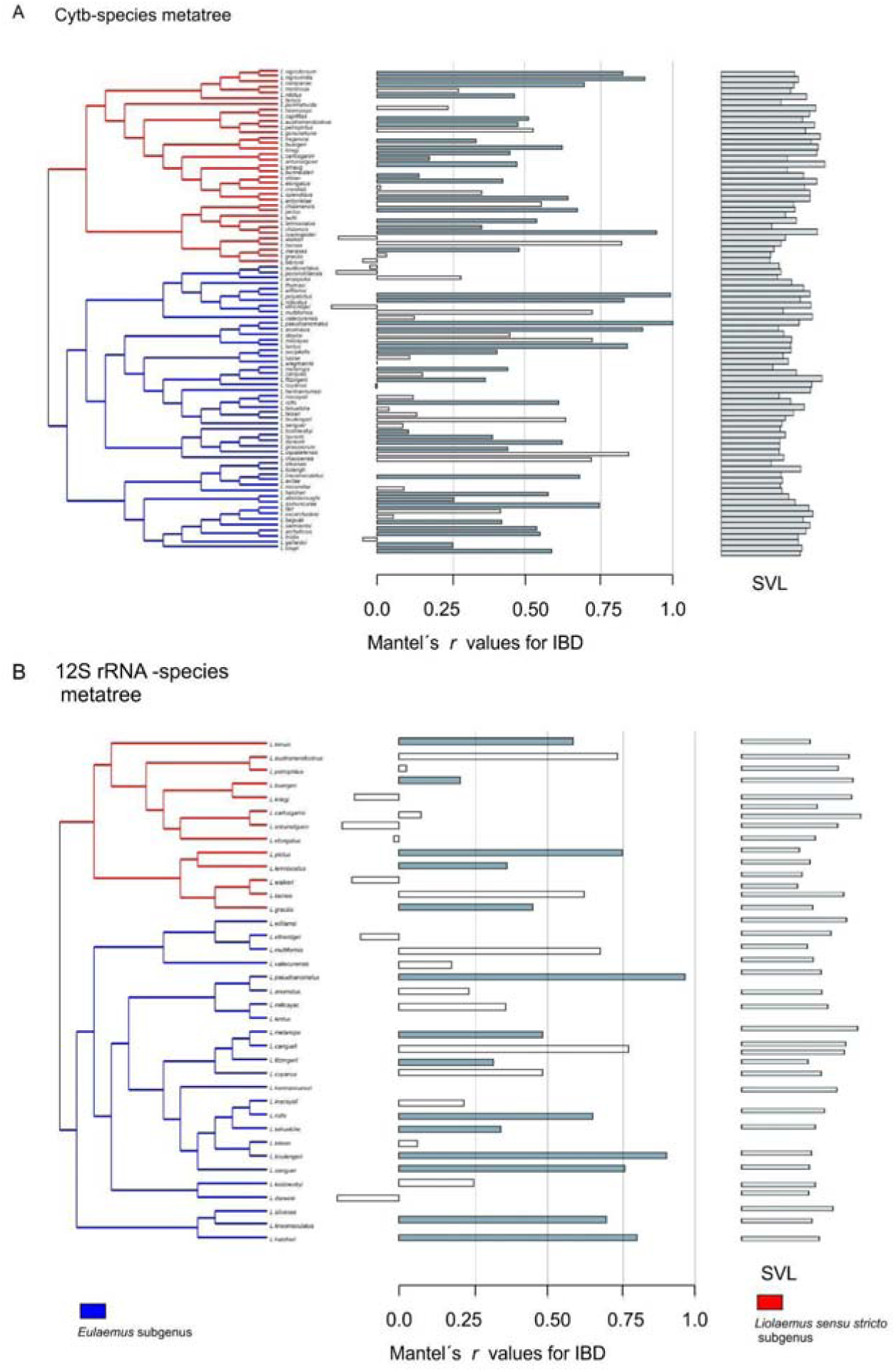
Meta-trees of *Liolaemus* lizards for (A) 86 species in the Cytb dataset and (B) 37 species in the 12S rRNA dataset. Branch colours indicate subgenera: red for *Liolaemus sensu stricto* and blue for *Eulaemus*. The middle panels show the correlation coefficient (*r*) from Mantel test of isolation by distance (IBD). Coloured bars represent statistically significant *r* values, while white bars indicate non-significant *r* values. The right panels display each species’ mean snout-vent length (SVL) (grey bars).

### Explanatory variables (Table 1): Number of specimens

We obtained the number of specimens (n) providing sequences for each species. We used this variable to test whether it constitutes one determinant of π-values (Table 1).

**Table 1:**
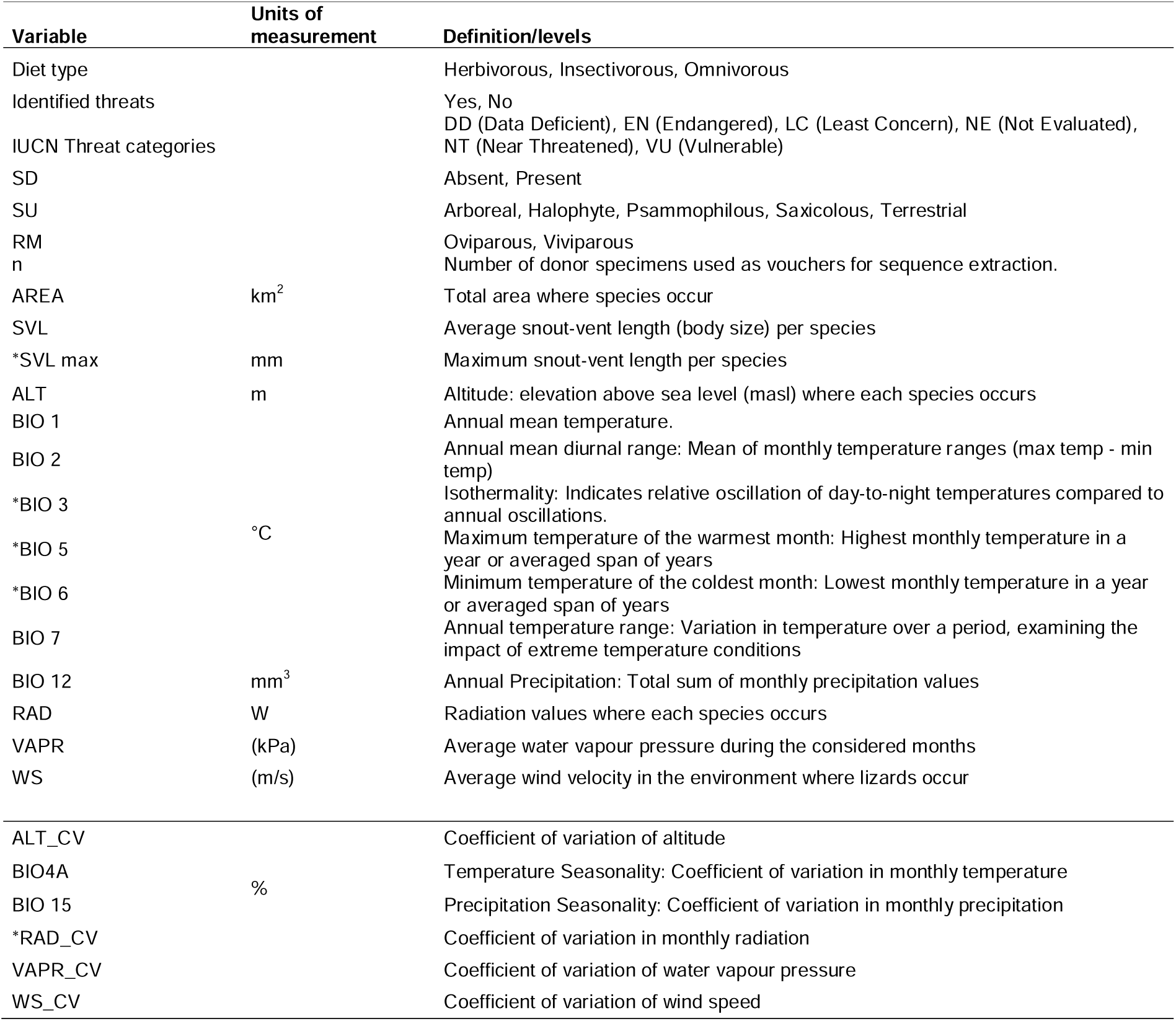
Variables used to examine potential determinants of genetic diversity in *Liolaemus* lizards. An asterisk (*) denotes variables that were highly correlated and therefore excluded from the analyses.

### Species traits

We compiled a suite of species-specific traits from the literature and our experience on the subject (Supplementary S4), including mean and maximum snout-vent length (SVL and SVL_max_), primary substrate use (Arboreal, Halophile —i.e., species that occur in salt terrain—, Psammophilous, Saxicolous, Terrestrial), diet (Herbivorous, Insectivorous, Omnivorous), reproductive mode (Oviparous, Viviparous), and presence of sexual dichromatism. Threat status (DD—Data Deficient—, EN— Endangered—, LC— Least Concern, NE—Not Evaluated—, NT—Near Threatened—, VU—Vulnerable—) and the presence of specific threats such as: agriculture expansion, energy production, natural, residential-commercial, viticulture settlements and transportation-service (Abdala et al., 2012, 2021; IUCN Red List 2024). We codified these as “Yes” or “No”.

### Geographic and abiotic data

We obtained geographic ranges for 77 species from the IUCN portal. For species lacking distribution data in the IUCN portal (e.g., *Liolaemus antonietae*, *L. anquapuka*, *L. audituvelatus*, *L. campanae*, *L. crandallii*, *L. meraxes*, *L. multiformis*, *L. nigrodorsum*, *L. splendidus*), we generated range shapefiles using QGis v3.36.0 (QGIS.org, 2024), incorporating data from herpetological collections (e.g., Colección de Herpetología IBIGEO, Instituto de Herpetología Fundación Miguel Lillo), species descriptions and relevant literature (Supplementaries S2, S4; Abdala et al., 2021). We calculated species area (km^2^) using QGis and included it as a predictor variable (Table 1).

To improve the representativeness, we generated 20 random points within each species’ geographic range, especially given the uneven sampling (Cytb Min =5, Max= 460; 12S rRNA Min = 5, Max = 54).

From the Worldclim v2.1 database (Flick and Hijmans, 2017), we extracted 11 abiotic variables (Table 1) at ∼19.87 km^2^ resolution. After checking for multicollinearity (Pearson’s *r* ≥ 0.60; *corrplot* package; Wei and Simko, 2021; Supplementary S5), we retained ALT (Altitude, masl), BIO1 (Annual mean temperature, °C), BIO2 (Annual mean diurnal range, °C), BIO7 (Annual temperature range, °C), BIO12 (Annual Precipitation, mm^3^), RAD (Radiation, W), VAPR (Water vapour pressure, kPa), and WS (Wind speed, m/s) (see; Table 1). We calculated the coefficient of variation (CV) for selected variables (ALT_CV, RAD_CV, VAPR_CV, and WS_CV) using data from 20 random points to assess environmental heterogeneity. Additionally, BIO4A (temperature CV) and BIO15 (precipitation CV) were included after excluding RAD_CV due to multicollinearity.

To visualise the geographic patterns of π-values, we mapped species ranges and plotted centroid coordinates associated with π. differentiating between reproductive modes and taxonomic groups (Fig. 5).

### Data analyses

#### Isolation by geographic distance (IBD)

For each species, we calculated intraspecific geographic distances (km^2^) using “distm” (*geosphere* package; Hijmans, 2022) from vouchers collection points. We compute intraspecific genetic distances, which provide one measure of genetic differences among populations, for Cytb and 12S rRNA genes (model = “K80”, Kimura 1980) with “dist.dna” function (*ape* package; Paradis and Schliep 2019). This function computes a matrix of pairwise distances from DNA sequences using a model of DNA evolution. Linear regressions between genetic and log_10_-transformed geographic distances allowed us obtaining by species-specific IBD slopes (β_IBD_). Non-significant β_IBD_ (*P* >0.05) were set to zero.

We then tested various explanatory variables (species area, number of specimens, substrate use, diet type, reproductive mode, ALT, SVL; Table 1) predicted β_IBD_ values. Before analysis, we tested for phylogenetic signal of β using Blomberǵs *K*, (where values closer to zero indicate weak phylogenetic patterns; Blomberg et al., 2003) and determined the best-fitting model of evolution explaining β_IBD_ values (Brownian Motion [BM], Early-Burst [EB], Ornstein-Uhlenbeck [OU]; Harmon et al., 2008; 2010) with “fitContinuous” (*geiger* package; Pennell et al., 2014; *ape* package Paradis and Schliep, 2019). Models selection was made following two criteria for chose those better-fitting models: the delta Akaike’s information criterion for small sample sizes (ΔAIC_C_) and Akaike’s weight (AIC_C*Wi*_), whose better models are indicated by ΔAIC_C_ values less than 2 and the highest AIC_C*Wi*_ value (Burnham and Anderson, 2002; Supplementary S6). β_IBD_ values (Cytb and 12S rRNA) presented weak phylogenetic patterns (Supplementary S6): β_IBD-Cytb_ *K*= 0.24, *P =* 0.214; β_IBD-12S rna_ *K*= 0.22, *P =* 0.69, and OU was the best evolutionary model for both (Supplementary S6). We fitted 126 PGLS models using the “gls” function (*nlme* package; Pinheiro and Bates, 2023), with an OU correlation structure using “corMartins” and selecting best fit models using ΔAIC_C_ and AIC_C*Wi*_ (Supplementary S6).

To assess IBD presence and its relationship with body size, we performed Mantel tests relating both data matrices, genetic distance with geographic distances using the “mantel.rtest” function (Pearson correlation, *ade4* package; Dray and Dufour, 2007) with 9999 permutations. Resulting correlation coefficients (*r*), *P*-values, and sample size (n) were recorded for each species. For species with non-significant *r* values (*P >* 0.05), we set *r* to = zero. We standardised and transformed *r* values into their effect sizes (*Zr r* and Var*Zr r*) using the “escalc” function (*metafor* package; Viechtbauer, 2010). We then employed a mixed-effects regression model using the “rma.mv” function from *metafor* to examine the relationship between *Zr r* and SVL. The SVL was log_10_-transformed, scaled, and fitted as a fixed effect. Phylogeny was incorporated as a random effect using a correlation squared matrix (A) derived from the phylogenetic tree (Fig. 2) using the “vcv” function (*ape* package; Paradis & Schliep 2019).

#### Genetic diversity

We examined whether π-values for Cytb and 12S rRNA were explained by morphological, life-history, and climatic factors as well as environmental heterogeneity (Table 1), thus we conducted multiple regression analyses using PGLS models (Supplementary S7). Phylogenetic signal of π-values for Cytb and 12S rRNA was assessed using Blomberǵs *K*, (Blomberg et al., 2003) and evolutionary models (Brownian Motion [BM], Early-Burst [EB], Ornstein-Uhlenbeck [OU]; Harmon et al., 2008; 2010) were tested to identify the best fit (Supplementary S6). Model selection was based on ΔAIC_C_ and AIC_C*Wi*.._Phylogenetic pattern was weak for π-Cytb (*K*= 0.28, *P =* 0.012) and stronger for π-12S rRNA (*K*= 1.25, *P=*0.001). The best-fitting evolutionary model for π-Cytb was the OU model (ΔAIC_C_ = 0; AIC_C*Wi*_ = 0.99), while for π-12S rRNA, the EB model was best (ΔAIC_C)_ = 0; AIC_C*Wi*_ = 0.68, negative *rate of change =* −0.54). We applied the “corMartins” (OU) for π-Cytb and “corBlomberg” (EB/ACDC model) functions for π-12S rRNA in PGLS analyses.

For numeric predictors (Table 1), we log_10_-transformed and scaled them to achieve a mean of zero and unit variance. We then fitted PGLS models using “gls” function (*nlme* package; Pinheiro and Bates, 2023), incorporating corMartins and corBlomberg correlation structures for π-Cytb and π-12S rRNA, respectively (Supplementary S6).

Due to missing data for some variables, we performed two sets of analyses: (i) using 12 numeric variables and a larger number of species (86 for Cytb and 37 12SrRNA), testing 4,095 models each; and (ii) incorporating categorical variables (e.g., reproductive mode, substrate use) with fewer species (60 for Cytb and 30 for 12SrRNA), testing 131,071 models each (Supplementary S7). Additionally, we tested combinations of five coefficient of variation in 31 models (Supplementary S7). Finally, we examined whether β_IBD_ was associated with π-values using similar PGLS model.

## Results

### DNA sequence retrieval

A search in the GenBank search using the term “*Liolaemus*” yielded 14,201 entries, of which 14,134 (corresponding to 238 described species and 96 undescribed species, cf. or sp.) contained *Liolaemus* species sequences (Supplementary S1). The most frequently studied genetic marker was Cytb (5,350 sequences), followed by the 12S rRNA (1,169 sequences). Other frequently represented markers included KIF24 (689 sequences), EXPH5 (525 sequences), C.most (467 sequences), and PRLR (461 sequences; Supplementary S1). Given the high representation of Cytb and 12S rRNA sequences, we focused our analyses on these two markers. Initially, Cytb data were available for 201 described and 62 undescribed *Liolaemus* species. After filtering, 3,293 sequences representing 86 described species remained (Figs. 1, 2A). For 12S rRNA, data were available for 170 described and 61 undescribed species. After filtering, 500 sequences from 37 described *Liolaemus* species remained (Figs. 1, 2B). In both cases, species with insufficient or non-geographically resolvable data were excluded from certain analyses.

### Isolation by Distance (IBD) patterns

#### Frequency of IBD across species

For Cytb IBD could be tested in 73 species (Fig.2A); as 13 species had samples from the same geographic locations, and therefore, we cannot calculate their geographic distance matrices. Among these, 41 species (56.16 %) exhibited significant Mantel correlations (*r*), indicating IBD patterns (Fig. 2A). Significant *r* values ranged from 0.1050 (*Liolaemus koslowskyi*) to 0.99 (*L. pseudoanomalus*), with a mean of 0.5541. For 12S rRNA, IBD could be tested in 33 species. Of these, 14 species (42.42%) showed significant IBD (*r* values from 0.205 in *L. buergeri* to 0.955 in *L. pseudoanomalus*; Fig. 2B).

#### Association of IBD with body size

For Cytb, including 73 species (setting non-significant *r* to 0) revealed no association between IBD (r) and SVL (Table 2). However, considering only those with significant *r* values (41 species) showed a significant negative association (Table 2; Fig.3), indicating that larger species tend to have lower IBD correlations. For 12S rRNA, no significant association between *r* and SVL was found, whether considering all 33 species or only those with significant IBD (Table 3).

**Fig.3:**
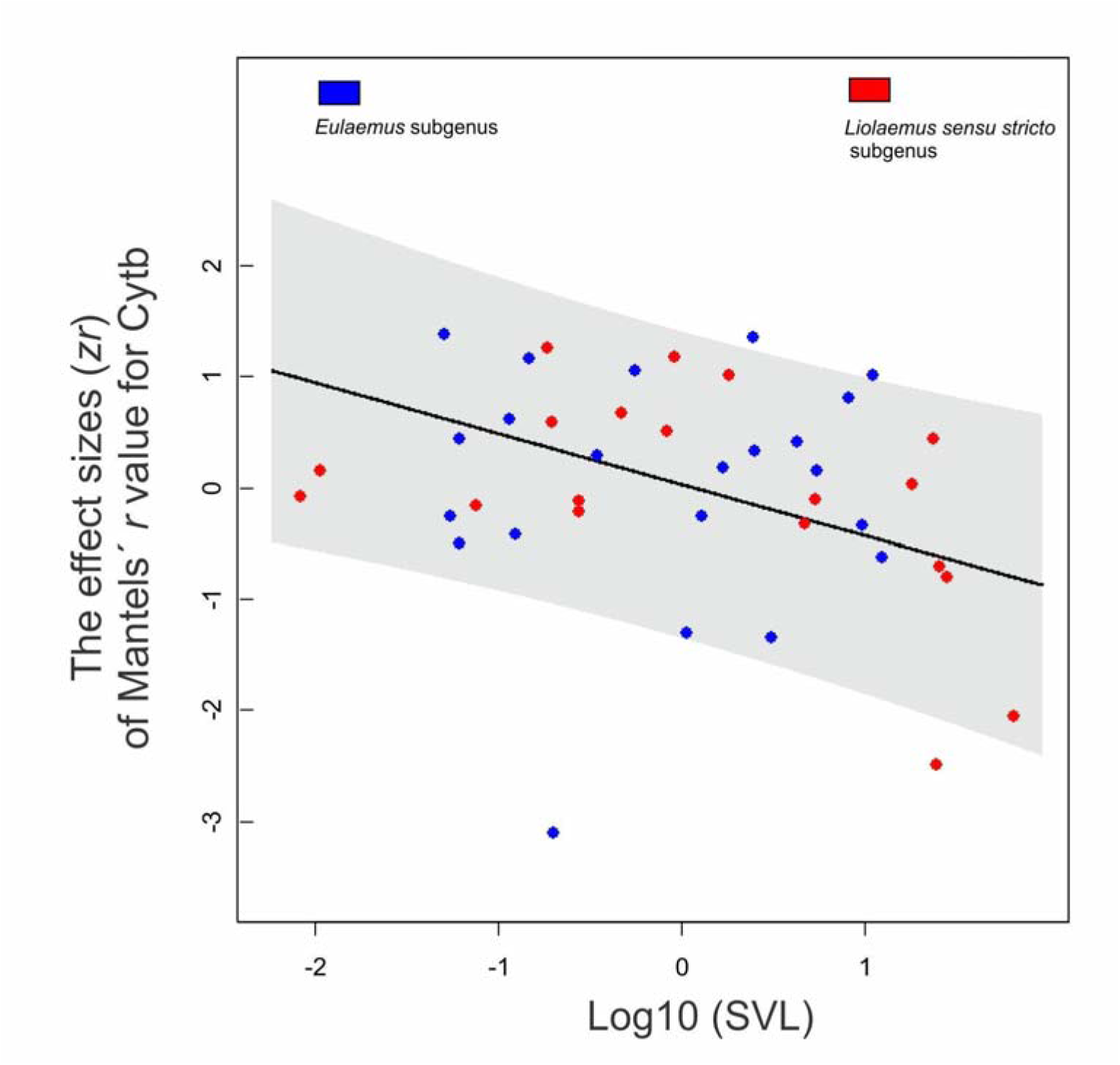
Negative association between the effect size of Mantel’s correlation coefficient (*r*) for isolation by distance (IBD) and snout-vent length (SVL). This analysis includes only species with significant IBD (*Zr r* > 0). Red and blue points represent species belonging to the *Liolaemus sensu stricto* and *Eulaemus* subgenera, respectively. The grey band indicates the 95 % confidence interval.

**Table 2:** Parameters for Cytb considering isolation by distance (IBD) and genetic diversity (π) analyses. The table shows models structure (Model); Number of species analysed (Nspp.), *F*-statistical (degree freedom), intercept, slope, coefficient of determination (*R*^2^) and *P*-values of models. In bold those results statistically significant (*P*<0.05).

| | Models | Nsp. | $F$ (df1, df2) | Intercept | Slope | $R^2$ | $P$ |
| --- | --- | --- | --- | --- | --- | --- | --- |
|  | IBD~SVL | 73 | 2.198 (1,71) | 0.544 | -0.074 | - | 0.142 |
|  | **IBD~SVL** | **41** | **8.04 (1,39)** | **0.028** | **-0.459** | **-** | **0.007** |
| IBD | $\beta_{\text{IBD-Cytb}} \sim \text{AREA}$ | **68** | **6.01 (1,66)** | **0.004** | **0.002** | **0.090** | **0.020** |
| | **Ovip**$\beta_{\text{IBD-Cytb}} \sim \text{AREA}$ | **29** | **3.92 (1,27)** | **0.003** | **0.003** | **0.130** | **0.049** |
| | Vivip$\beta_{\text{IBD-Cytb}} \sim \text{AREA}$ | 39 | 1.78 (1,37) | 0.003 | 0.001 | 0.050 | 0.190 |
| | **Eul.**$\beta_{\text{IBD-Cytb}} \sim \text{AREA}$ | **43** | **6.94 (1,41)** | **0.003** | **0.002** | **0.140** | **0.012** |
| | Lio.ss$\beta_{\text{IBD-Cytb}} \sim \text{AREA}$ | 25 | 1.34 (1,23) | 0.005 | 0.002 | 0.055 | 0.260 |
| | $\pi \sim \text{SVL}$ | **85** | **3.96 (1,83)** | **0.019** | **-0.004** | **0.050** | **0.049** |
| $\pi$ | **Ovip.**$\pi \sim \text{SVL}$ | **31** | **4.36 (1,29)** | **0.022** | **-0.007** | **-0.130** | **0.045** |
| | **Vivip.**$\pi \sim \text{SVL}$ | **49** | **5.67 (1,47)** | **0.020** | **-0.007** | **0.110** | **0.020** |
| | Eul.$\pi \sim \text{SVL}$ | 50 | 0.047 (1,48) | 0.018 | 0.001 | -0.001 | 0.820 |
| | **Lio.ss**$\pi \sim \text{SVL}$ | **35** | **9.83 (1,33)** | **0.023** | **-0.010** | **0.220** | **<0.01** |
| | $\pi \sim \text{BIO7}$ | **85** | **10.26 (1,83)** | **0.019** | **-0.008** | **0.110** | **< 0.01** |
| | Ovip.$\pi \sim \text{BIO7}$ | 31 | 0.27 (1,29) | 0.025 | -0.002 | -0.770 | 0.600 |
| | **Vivip.**$\pi \sim \text{BIO7}$ | **49** | **20.54 (1,47)** | **0.014** | **-0.012** | **0.300** | **< 0.001** |
| | Eul.$\pi \sim \text{BIO7}$ | 50 | 3.01 (1,48) | 0.017 | -0.004 | 0.050 | 0.090 |
| | **Lio.ss**$\pi \sim \text{BIO7}$ | **35** | **13.42 (1, 33)** | **0.023** | **-0.015** | **0.320** | **< 0.01** |
| | $\pi \sim \text{BIO4A}$ | **85** | **15.116 (1, 83)** | **0.019** | **-0.007** | **0.130** | **< 0.01** |
| | Ovip.$\pi \sim \text{BIO4A}$ | 31 | 0.16 (1, 29) | 0.025 | -0.003 | -0.008 | 0.690 |
| | **Vivip.**$\pi \sim \text{BIO4A}$ | **49** | **15.12 (1, 47)** | **0.016** | **-0.008** | **0.240** | **< 0.001** |
| | **Eul.**$\pi \sim \text{BIO4A}$ | **50** | **8.01 (1, 48)** | **0.017** | **-0.006** | **0.150** | **< 0.01** |
| | **Lio.ss**$\pi \sim \text{BIO4A}$ | **35** | **4.94 (1, 33)** | **0.023** | **-0.012** | **0.160** | **0.030** |

**Table 3:**
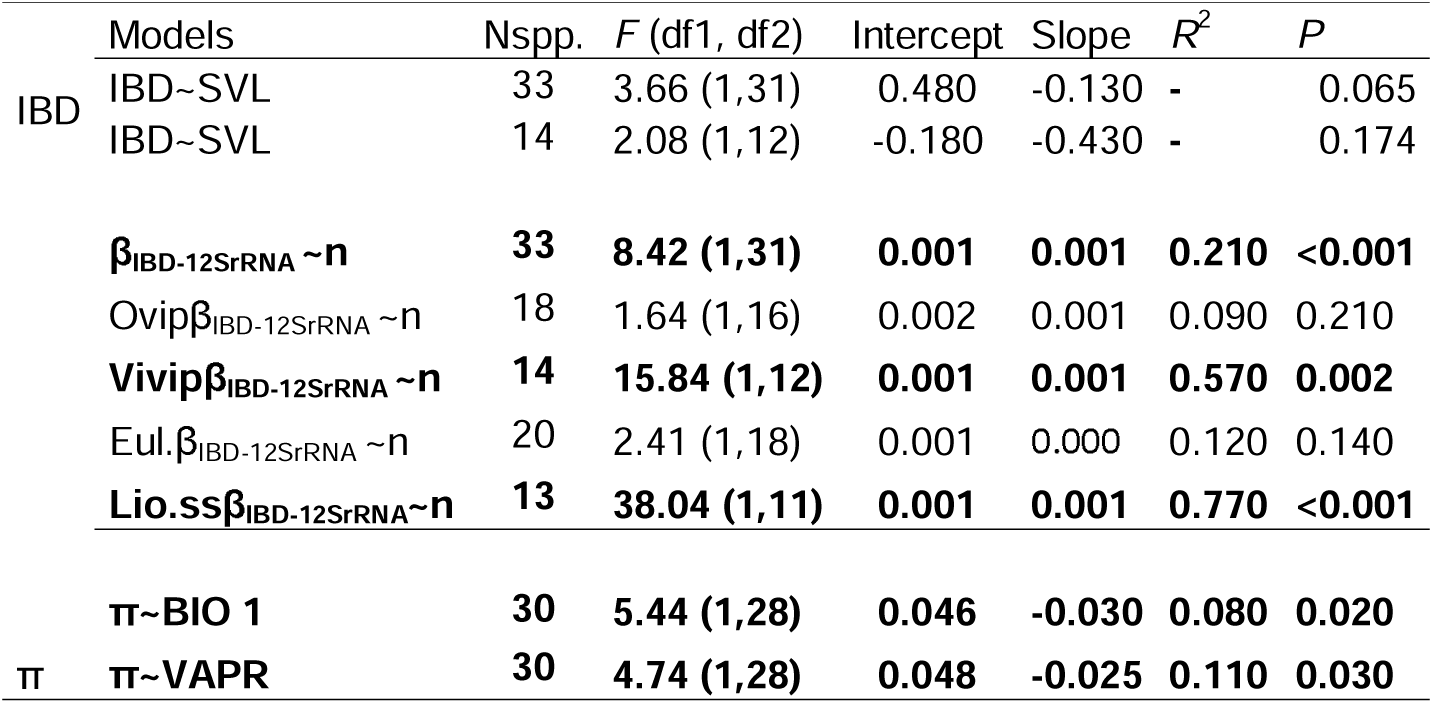
Parameters for 12SrRNA considering isolation by distance (IBD) and genetic diversity (π) analyses. The table shows models structure (Model); Number of species analysed (Nspp.), *F*-statistical (degree freedom), intercept, slope, coefficient of determination (*R*^2^) and *P*-values of models. In bold those results statistically significant (*P*<0.05).

#### Determinants of the IBD slope

For Cytb, β_IBD-Cytb_ could be calculated for the same 73 species. Among these, 57 species (78.08 %) exhibited significant β_IBD-Cytb_ (Supplementary S8). Values ranged from 0 to 0.024 (*Liolaemus gununakuna*). The best-fitting model (β_IBD-Cytb_ ∼AREA; Supplementary S7) identified a positive association of β_IBD-Cytb_ with geographic range size (Table 2; Fig. 4). This relationship was significant for oviparous species and for *Eulaemus* subgenus (Table 2), but not for viviparous species or *Liolaemus sensu stricto* (Table 2; Figs. 4A-B).

**Fig.4:**
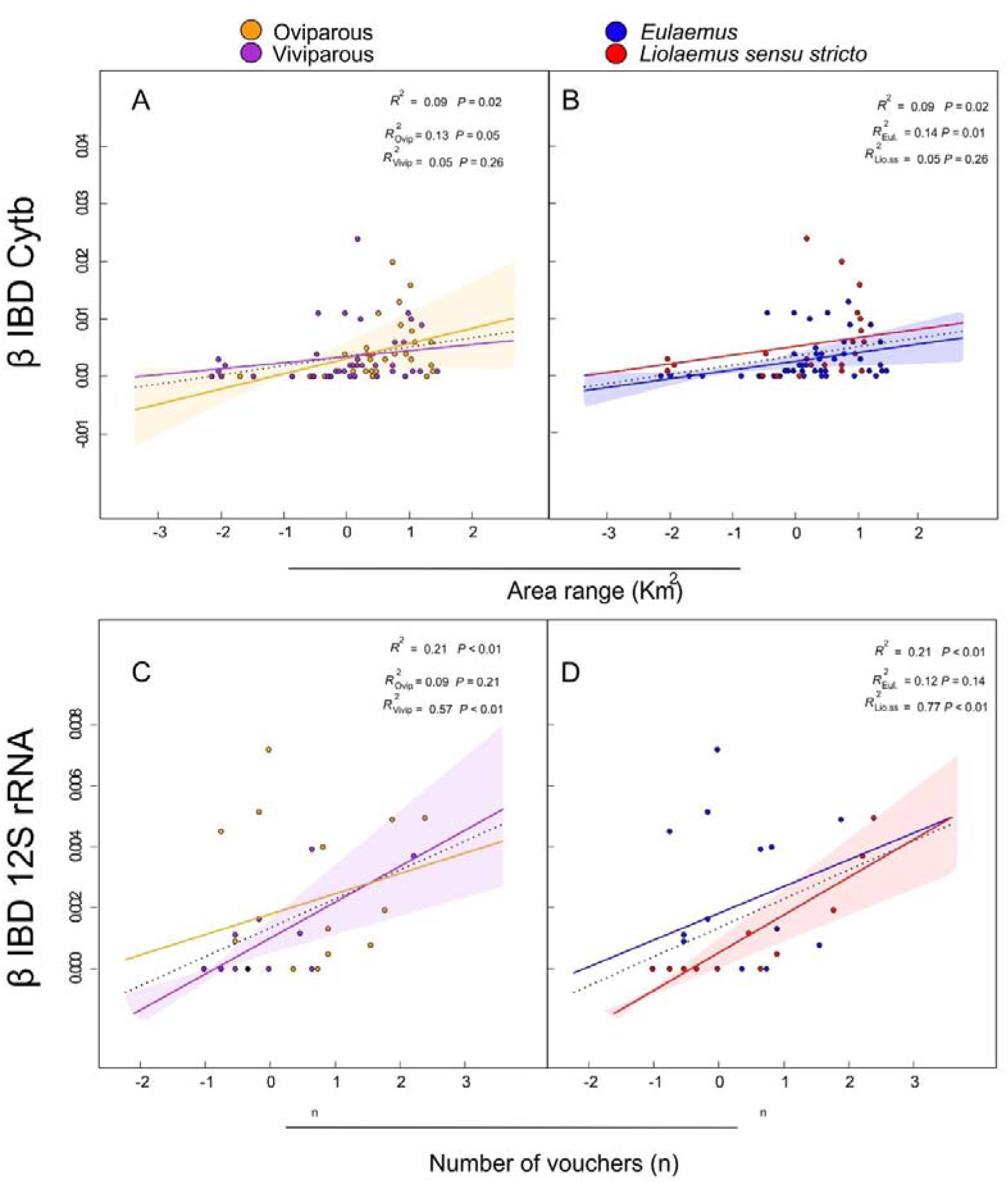
Positive associations of the isolation by distance (IBD) slope (β) in Cytb (A-B) and 12S rRNA (C-D). For Cytb, β is positively correlated with species’ geographic range size, whereas for 12S rRNA (C-D), β is positively correlated with sampling size. The left panels show comparisons by reproductive mode (orange: oviparous, purple: viviparous), and the right panels show comparisons by sub genus (red: *Liolaemus sensu stricto*, blue: *Eulaemus*).

For 12S rRNA, β_IBD-12S rRNA_ values were significant in 16 of the 33 tested species (48.49%; Supplementary S8). Values ranged from 0 (17 species) to 0.0072 (*Liolaemus boulengeri*) with a mean of 0.0014. Best fitted model (β_IBD-12S RNA_ ∼n; Supplementary S7) exhibited a positive association of the number of specimens with β_IBD-12S rRNA_ (Table 3). This relationship was significant for viviparous species and *Liolaemus sensu stricto* (Table 3; Fig. 4D), but not for oviparous species or *Eulaemus* (Table 3; Fig. 4D).

#### Genetic diversity (π) estimates

For Cytb, π values across the 86 species ranged from 0 (*Liolaemus hermannunezi*) to 0.225 (*L. cuyanus*), mean = 0.021. Because *L. cuyanus* contributed disproportionately large to variance (57.64 %), it was excluded. Without it, 85 species showed π-Cytb values ranged from 0 to 0.081 (*L. cyanogaster*), mean = 0.019. Geographically, species were distributed broader latitudinally (from 11 °N *Liolaemus walkeri* to ∼ 51 °S *L. sarmientoi*; Figs. 5A-B) than longitudinally (42 °E *L. lutzae* to ∼ 75 °W *L. walkeri*). Higher values of π-Cytb appeared in Andean and southern regions of Argentina and Chile (Figs.5A-B). Viviparous species and *Eulaemus* subgenus covered greater latitudes, and π values decreased with latitude increases (Vivip. PGLS *F* = 14.68 _(1,47)_, *P* = 0.03, slope = −0.0006, intercept = 0.04, *R*^2^ = 0.13; *Eulaem*. PGLS *F* = 15.42 _(1,48)_, *P* = 0.02, slope = −0.0005, intercept = 0.03, *R*^2^ = 0.10). While longitude geographic increases favoured π in viviparous and *Liolaemus sensu stricto* species (Vivip.PGLS *F* = 14.77 _(1,47)_, *P* = 0.03, slope = 0.0035, intercept = −0.232, *R*^2^ = 0.13; *Liol.ss*. PGLS *F* = 110.83 _(1,33)_, *P* < 0.01, slope = 0.006, intercept = −0.41, *R*^2^ = −0.599).

**Fig.5:**
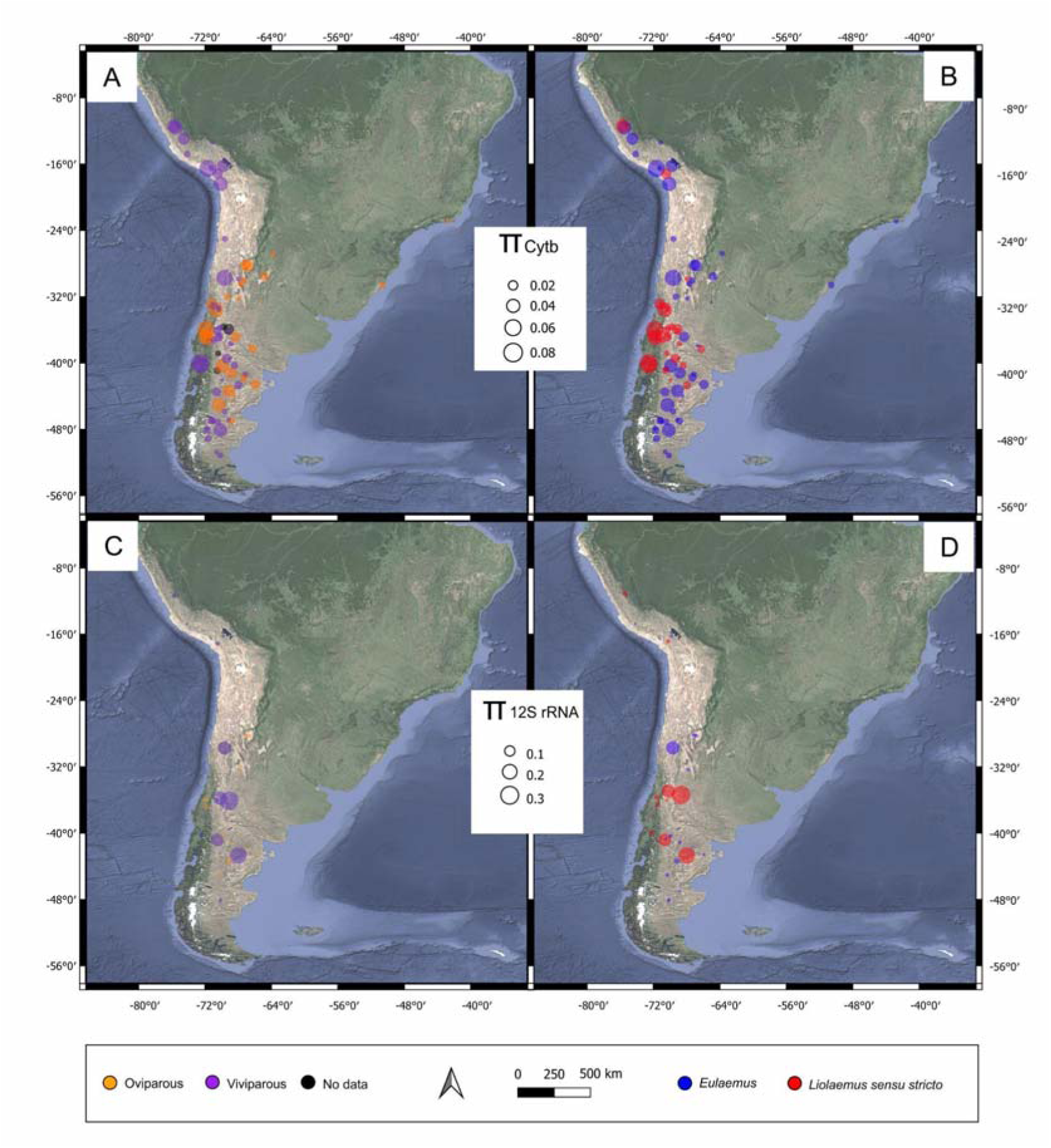
Geographic centroids of *Liolaemus* species displaying nucleotide diversity (π), values for Cytb (A-B) and 12SrRNA (C-D). In each panel circle sizes within insert box represent π values. The left panels show species by reproductive mode (orange: oviparous, purple: viviparous), while the right panels show species by sub genus (red: *Liolaemus sensu stricto*, blue: *Eulaemus*).

For 12S rRNA, π values ranged 0 (*Liolaemus hermannunezi*, *L. lentus*) to 0.32 (*L. austromendocinus*), mean = 0.0364. Species ranged broadly from 11 °N (*L. walkeri*) to 48 °S (*L. hatcheri*; Figs. 5C-D) and from 65 °E (*L. melanops*) to 75 °W (*L. walkeri*; Figs.5C-D). The highest π-12S rRNA values were found in southern Argentina, particularly within four *Liolaemus sensu stricto* species (*L. austromendocinus*, *L. buergeri*, *L. kriegi*, *L. petrophilus*; Fig. 5D). π values did no evidenced significant association with latitude and longitude.

#### Determinants of genetic diversity (π)

Morphological and life-history predictors. When considering all explanatory variables (n = 60 species), SVL emerged as the primary predictor (PGLS *F* _(1, 58)_ = 7.55, *P* < 0.01, slope = −0.0069, *R*^2^ = 0.09; Supplementary S7). The SVL was also statistically significant for oviparous species (*F* _(1, 26)_ = 4.305, *P* < 0.05, slope = −0.012), but not for viviparous species. Pooling data (85 species) also showed a significant and negative association with SVL (Table 2), this pattern also was observed in both reproductive mode and for species belonging to *Liolaemus sensu stricto* (Fig. 6A-B Table 2).

**Fig. 6:**
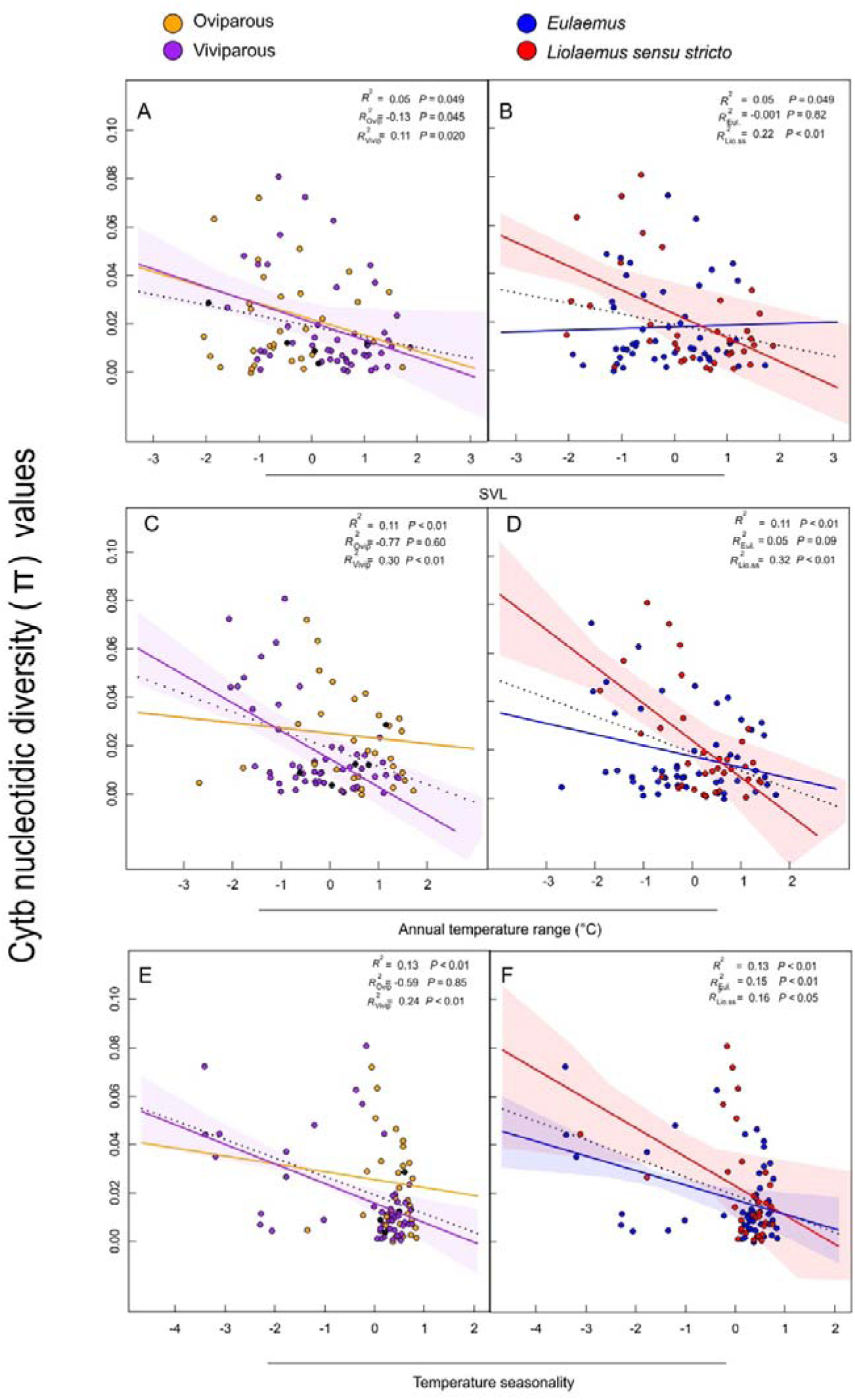
Negative associations of Cytb nucleotide diversity (π) with (A)-annual temperature range, (B)-snout-vent length and (C), - temperature coefficient of variation. The left panels show reproductive mode (orange: oviparous, purple: viviparous), while the right panels show sub genus groupings (red: *Liolaemus sensu stricto*, blue: *Eulaemus*).

For π-12S rRNA, models including numeric and categorical variables (30 species), five models were identified as better-fitting (Supplementary S7). Only models with annual mean temperature (BIO 1) and water vapour pressure (VAPR) showed a negative and significant with π-12S rRNA (Table 3). The models did not yield significant results considering both reproductive mode or subgenus.

Environmental and climatic predictors. For π-Cytb, using only numeric variables (85 species), annual temperature range (BIO 7) was the primary and negative predictor (Table 2; Supplementary S7). This predictor was significant for viviparous species and *Liolaemus sensu stricto* (Table 2; Fig. 6C-D) but not for oviparous species or *Eulaemus* (Table 2). Considering environmental heterogeneity (obtained from different climatic coefficient of variation), temperature seasonality (BIO4A) emerged as the best and negative predictor of π-Cytb (Table 2, Figs. 6E-F, Supplementary S7), again significant mainly for viviparous species (Table 2; Fig. 6E) and both subgenera (Table 2). For π-12S rRNA, five models were best fitted (Supplementary S7) but these models did not yield significant results.

#### Relationship between β_IBD_ and π

For Cytb, there was a significant association between π and β_IBD_ values (PGLS *F* _(1, 70)_ = 26.80, *P* < 0.001, slope = 1.776, *R*^2^ = 0.283; Fig. 7), indicating that species with stronger IBD patterns also tended to have higher nucleotide diversity. In contrast, non-significant association was found between π-12S rRNA and β_IBD-12S rRNA_ values (PGLS *F* _(1, 31)_ = 0.25, *P* = 0.62).

**Fig.7:**
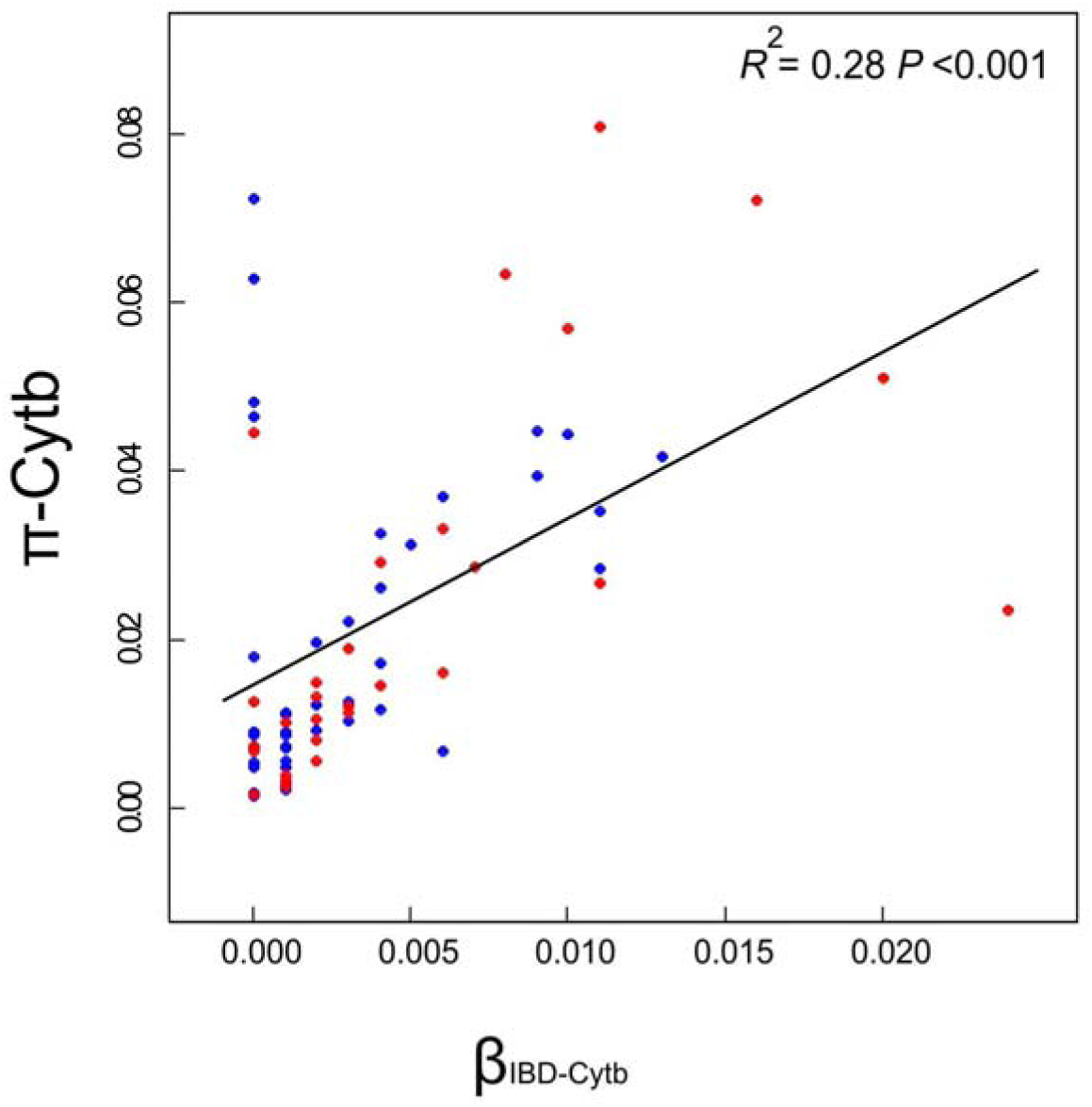
Positive association between the isolation-by-distance (IBD) slope (β) and Cytb nucleotidic diversity (π) in *Liolaemus* lizards. Red and blue points represent species belonging to the *Liolaemus sensu stricto* and *Eulaemus* subgenera, respectively.

## Discussion

Our study provides new insights into genetic diversity and its environmental (climate and heterogeneity) and phylogenetic determinants in the highly diverse genus of *Liolaemus* lizards. Two patterns emerged, first, isolation by distance (IBD) was widespread, particularly for the Cytb gene, where larger species ranges and body size explained IBD patterns, while the sampling effort (number of individuals) influence IBD patterns in 12S rRNA gene. Second, genetic diversity (π), was shaped by distinct factors depending on the gene studied, thermal variation (annual temperature range)/heterogeneity (temperature coefficient of variation) and IBD influenced π-Cytb, while phylogenetic relationships among species explained π-12S rRNA. These patterns further examined through reproductive modes (oviparous and viviparous species) and taxonomic groupings (*Eulaemus* and *Liolaemus sensu stricto*) highlight the complex interplay of evolutionary and ecological factors that shape genetic diversity in *Liolaemus*.

### Isolation by distance (IBD)

IBD was common across *Liolaemus* with 56.16% for Cytb and 42.42% for 12S rRNA of tested species showing significant IBD. These results align with previous studies suggesting that geographic isolation, topographic barriers, and allopatric fragmentation, often linked to Andean uplift, have significantly promoted diversity in Liolaemidae family (Esquerré et al., 2019). IBD is variable in *Liolaemus* lizards, with clear, absent or mixed IBD patterns (e.g., Morando et al., 2007; Vera-Escalona, 2012; Villamil et al., 2017; Cab-Sulub and Álvarez-Castañeda, 2022; Sánchez et al., 2024), similar with to the observed for the Squamate order (Jenkins et al., 2010).

For the Cytb, larger body size was associated with weaker IBD, possibly because larger species tend to have greater home ranges and dispersal capabilities (Perry and Garland Jr, 2002), which could eventually attenuate the impact of geographic isolation on gene flow. This association was no detected for 12S rRNA, likely due to the smaller sample size for this gene (only 14 species presented significant IBD), which highlight the need for future studies including more species and markers.

### Determinants of β_IBD_

Factors explaining β_IBD_ varied with the genetic marker. In the case of the Cytb gene, species with larger geographic ranges had steeper IBD slopes, suggesting that widely distributed species experience greater genetic differentiation as geographic distance increases. Similar patterns have been observed in geckos and in North American squamates (Larkin et al., 2023; Lau et al., 2024). Conversely, for 12S rRNA, the best predictor of β_IBD_ was the “number of specimens” suggesting a sampling effect. High sample size can revel genetic differentiation within species (Hague and Routman, 2016; but see Rutherford et al., 2019).

Patterns of β_IBD-Cytb_ differed between reproductive modes and taxonomic groups. Oviparous species often extend southward without being restricted to the Andes (it is possible that the Andes act as a barrier to expand species ranges, Esquerré et al., 2019), potentially increasing the species-areas and favouring gene flow disruption. While *Eulaemus* species have a broader longitudinal distribution, which may enhance IBD effects. In contrast for β_IBD-12S rRNA_, where viviparous species and species belonging to *Liolaemus sensu stricto* showed that sample size drive β. We observed that four viviparous species from Argentinian Patagonian, (a region with high species diversity; Avila et al., 2020; Morando et al., 2020; Abdala et al., 2021), exhibited higher π in 12SrRNA gene. Then, it possible that in this environment with high species richness and potential competition, populations that are larger and genetically more diverse are likely to be more successful than smaller populations with lower genetic diversity. Also, Patagonian species with larger and extended populations better buffer effect temperature variation (Kubisch et al., 2023).

We also found that β_IBD-Cytb_ was positively associated with π-Cytb (*R*^2^ =0.28). Geographic distance similarly influences genetic differentiation in various organism including lizards (Fonseca et al., 2024; McGreevy Jr et al., 2024) and other species (Li et al., 2021; Baccaro et al., 2024). This widespread pattern suggests that geographic distance is a fundamental driver of genetic diversity across different taxa.

### Environmental determinants of genetic diversity

For π-Cytb, an Ornstein–Uhlenbeck (OU) evolutionary model was best, similar to Larkin et al. (2023 in squamate lizards at least), indicating selective pressures may drive Cytb diversity toward one or more evolutionary optima (Martins and Hansell, 1997; Harmon et al., 2008; 2010). Although, it is important to mention that factors like low species sampling or data errors could lead to overestimating an OU process in certain cases (Cooper et al., 2015); nevertheless, our number of species is quite important, 85 species (almost one-third of the total species in the genus).

Environmental factors, specifically thermal variation (annual temperature range—BIO 7) and heterogeneity (temperature coefficient of variation— BIO4A) emerged as main determinants of π-Cytb. The effect was stronger in viviparous species, from colder, more hostile Andean and austral habitats, where efficient thermoregulation and higher respiratory rates may have shaped Cytb evolution (Clavijo-Baquet et al., 2022; Cruz et al., 2022; Penman et al., 2022). Similar environmental determinants of genetic diversity have been reported in amphibians and in other taxa (Miraldo et al., 2016; Amador et al., 2024; Sexton et al., 2013), reinforcing those climatic and environmental gradients are key shapers of intraspecific genetic variation in ectotherms. Finally, we observed that for Cytb, latitude has been negatively correlated with genetic diversity, especially in viviparous species. This a common and broad pattern observed in several taxa, (Brüniche-Olsen et al., 2018; Pelletier and Carstens, 2018; Smith et al., 2017; Amador et al., 2024).

### Phylogenetic determinants

In contrast to Cytb, phylogenetic relationships influenced more strongly π-12S rRNA values. The best-fitting model for π-12 S rRNA was an Early-Burst (EB) model, suggesting that π-12 S rRNA genetic diversity patterns could have been established during early diversification events within *Liolaemus*, particularly during the split into *Eulaemus* and *Liolaemus sensu stricto* subgenera (Esquerré et al., 2019). The strong phylogenetic signal in π-12S rRNA suggests that closely related species share more similar genetic diversity patterns due to shared ancestry rather than independent environmental or ecological influences (Romiguier et al., 2014). However, we must be cautious as we counted with a low number of species in the case of 12S rRNA (37 species) and there could be no balanced taxonomic representativeness, strengthening phylogenetic patterns in those clades more represented. Although we observed that environmental variables like water vapour pressure and annual mean temperature were identified as predictors in a reduced dataset (30 species), these results were influenced by a single species (*Liolaemus vallecurensis*) with moderate high π-12 S rRNA value (0.16) and lowest humidity and mean temperature. Thus, the phylogenetic context likely offers a more robust explanation for π-12S rRNA patterns than local environmental factors.

Hybridization events in *Liolaemus* further compound the complexity of phylogenetic determinants. The “Promiscuous ones and the successful Generalist Hypothesis” (Morando et al., 2020), proposes that hybridization promotes the diversity of the *Liolaemus* genus with multiple species with a possible hybrid origin (details in Morando et al., 2020). Then, despite morphological differentiation in species of this genus, genetic divergence in some groups is scarcely observed or hybridization cases among close sibling species are frequent (Medina et al., 2015; 2017; Olave et al., 2011, 2018; Araya-Donoso et al., 2019; Grummer et al., 2021; Sánchez et al., 2024). For example, *L. parthenos* displays mitochondrial introgression (Abdala et al., 2016), and asymmetrical mtDNA introgressions occur in *L. bibronii* and *L. gracilis* species (Olave et al., 2011). Such cases may homogenize genetic diversity among closely related species, reinforcing the importance of phylogenetic context and introgressive hybridization in shaping genetic diversity.

### Conclusions

Our results indicate that ecological, and environmental factors, as well as phylogenetic history, shape patterns of genetic diversity in *Liolaemus* lizards. For Cytb, geographic isolation (IBD) and thermal environmental factors appear to drive selective evolution of intraspecific genetic diversity, while, for 12S rRNA, shared evolutionary history and phylogenetic relatedness are more influential. These contrasting patterns underscore the importance of considering multiple genetic markers, environmental gradients, reproductive modes, and phylogenetic relationships when studying genetic diversity in diverse radiations like *Liolaemus*.

By integrating ecological and evolutionary contexts, our findings provide insights into how genetic diversity may be maintained and structured in this species-rich genus. Future research incorporating additional life-history traits, ecological niches, and genomic data will further elucidate the multifaceted processes influencing genetic diversity in *Liolaemus* and other diverse lineages.

## Supplementary data

https://data.mendeley.com/datasets/6r76dfh4wv/2

## Acknowledgments

We thank C. S. Abdala and A. S. Quinteros for their assistance with some species determinations and discussions on taxonomic issues and distributional ranges. The authors are career researchers at the Consejo Nacional de Investigaciones Científicas y Técnicas (CONICET). This study was partially funded by CONICET (PIP 2022-2024) awarded to JJM, MRR-M, and Lucía Sommaro.

